# found: Inferring cell-level perturbation from structured label noise in single-cell data

**DOI:** 10.64898/2026.04.10.717768

**Authors:** Elia Afanasiev, Aleksandrina Goeva

**Affiliations:** Donnelly Centre for Cellular and Biomolecular Research, University of Toronto, Toronto, ON M5S 3E1, Canada

**Keywords:** perturbation inference, single-cell RNA-seq, label refinement, machine learning, computational biology

## Abstract

Recent work by Goeva *et al*. introduced HiDDEN, a method for refining batch-level labels to infer cell-level perturbation without prior knowledge of affected populations, addressing the mismatch between sample-level labels and heterogeneous perturbation effects across cells.

Here, we present `found`, a Python and R implementation of HiDDEN, supporting pipeline customization, by-factor grouping, hyperparameter selection, and visualization. Through benchmarking across diverse datasets, we show that performance depends strongly on modeling choices, particularly regression, grouping, and embedding dimensionality. `found` provides a practical, flexible, and accessible framework for robust cell-level perturbation analysis.

## Background

Single-cell -omics technologies enable the study of biological systems at cellular resolution, yet subtle and heterogeneous perturbation signals are often obscured by dominant sources of variation, such as cell type, cell cycle, and technical/experimental noise. In case-control studies, condition labels are assigned at the sample level and propagated to all constituent cells. When perturbation effects are strong or widespread, affected cells may separate clearly; however, in more common intermediate regimes – where only a subset of cells is affected – the signal can remain diluted and difficult to detect, particularly in rarer cell populations.

Existing approaches typically address this mismatch through experimental enrichment (e.g. cell sorting via flow cytometry) or *in silico* filtering. However, these strategies require prior knowledge of affected populations or sufficiently strong signals, and are therefore limited in exploratory settings where perturbation effects are weak or heterogeneous.

HiDDEN addresses this challenge by inferring cell-level perturbation signal from sample-level labels affected by structured label noise, producing both continuous scores and refined binary labels for each cell [1]. This reframes case-control analysis as a latent variable problem at the level of individual cells.

In practice, applying HiDDEN requires selecting among modeling choices at each stage of the pipeline, including embedding, scoring, discretization, and grouping strategies, with performance that can vary substantially across datasets.

Here, we present `found`, a Python and R implementation of HiDDEN designed to support flexible pipeline composition so the modeling framework can be customized to the dataset of interest, systematic evaluation which facilitates a streamlined comparison of modeling choices, and reproducible analysis across diverse single-cell workflows.

## Results and Discussion

### Overview of HiDDEN and existing solutions

HiDDEN was previously shown to outperform existing approaches for detecting perturbation-affected sub-populations in case-control single-cell data [1]. HiDDEN is best understood not as a fixed method, but as a family of pipelines whose behavior depends on modeling choices at each stage.

HiDDEN begins by processing the high-dimensional -omics dataset to generate lower-dimensional embeddings (fig. 1a) [1]. While this step is theoretically optional, it is effectively required in practice. First, regression methods struggle with co-linear features, which are common in high-dimensional -omics datasets. Second, raw count matrices contain confounding signals that should be controlled for, such as size-factor-related technical noise or per-sample biologically irrelevant batch effects [2], [3]. The embedding step also introduces a key hyperparameter: the dimensionality of the resulting embedding space, denoted *k* (fig. 1a).

**Figure 1:**
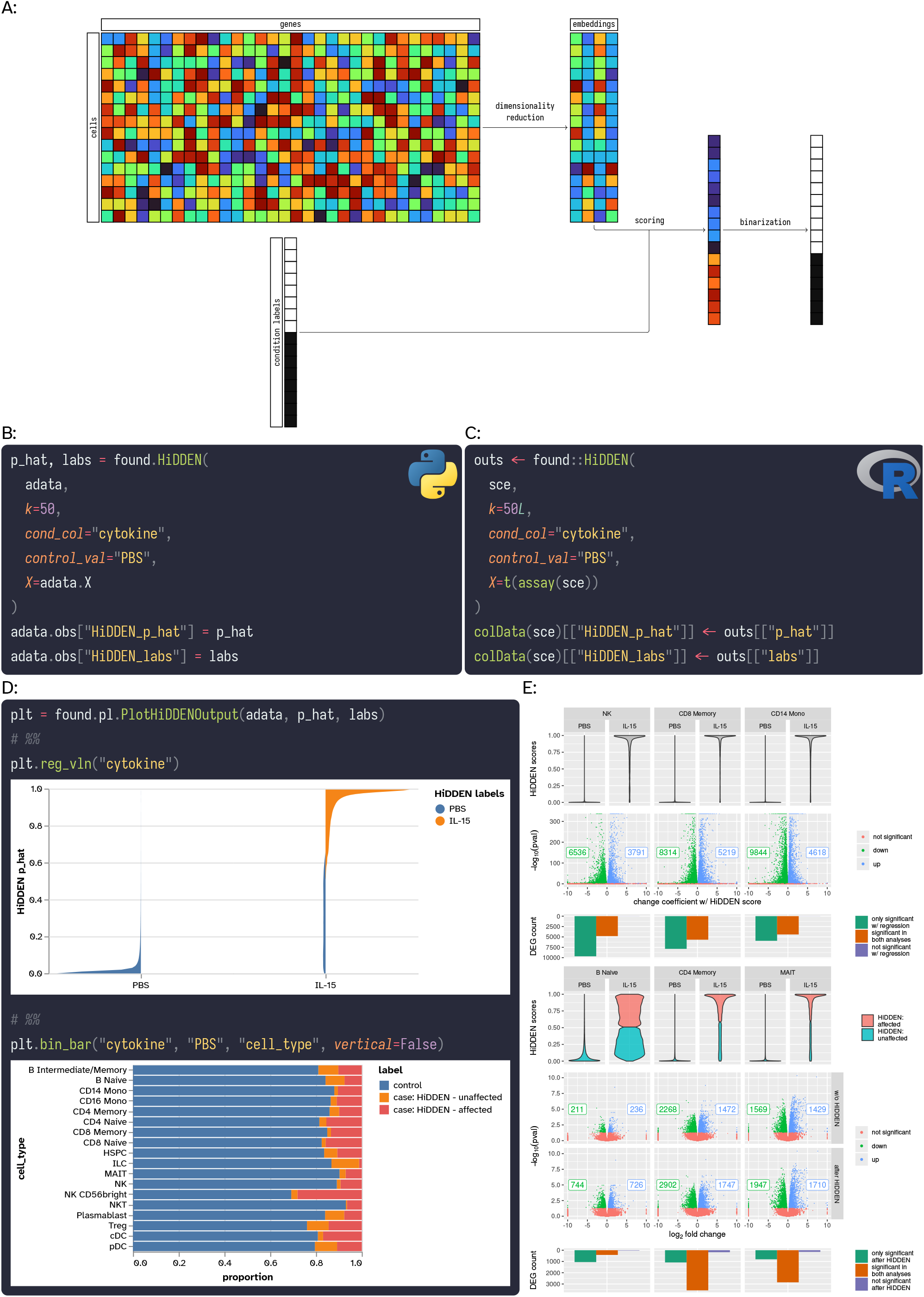
HiDDEN/foundoverview and demonstration. A: Schematic of the HiDDEN pipeline, including embedding, continuous scoring, and discretization steps. B: Example usage of the `found` library in Python. C: Example usage of the `found` library in R. D: Example usage of visualization utilities from the `found.pl` module in Python. E: Evaluation of HiDDEN outputs on IL-15-stimulated PBMC data (PBS as control). Top rows: Violin plots showing 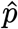 (y-axis) in PBS and IL-15 stimulated cells (x-axis), across cell types in different columns (natural killer (NK) cells, CD8 memory cells, and CD14 Monocytes (mono) in left-to-right order). Middle rows: Gene-level statistics showing association with 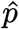 (x-axis) and differential expression − log_10_(pval) values (y-axis) across conditions. Bottom rows: Comparison of differential expression results using (i) continuous 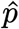, (ii) original batch-level labels, and (iii) HiDDEN-refined labels. 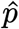 : inferred cell-level perturbation score. HiDDEN labels: binary labels derived from 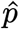.

Following embedding, the batch-level labels and embeddings are jointly used via a predictive model to assign each cell a continuous perturbation score, denoted 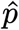 (fig. 1a) [1]. Based on the distribution of 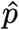, a binarization step can be applied to refine batch-level labels, relabeling case cells with a low 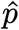 as “unaffected” (fig. 1a) [1]. Alternatively, continuous perturbation scores can be used directly in downstream analyses when a gradient interpretation is more appropriate.

This formulation is intentionally abstract, allowing for different methods to be used at each step. In the original HiDDEN work, Goeva *et al*. evaluated several choices and recommended PCA for embedding generation, a logistic regression for 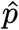 generation, and 2-means clustering for the optional discretization step [1]. Additionally, two heuristics were proposed for selecting *k* without prior knowledge of the dataset [1].

### found-HiDDEN implementation

Here, we present `found`, a comprehensive implementation of the HiDDEN framework designed to enable its practical and reproducible use. Rather than providing a fixed pipeline, `found` exposes HiDDEN as a flexible modeling framework, supporting multiple choices for embedding, continuous scoring, and discretization, along with tools for hyperparameter selection and output evaluation. We additionally implement extended routines for dimensionality reduction and regression not present in the original implementation. We further extend hyperparameter selection and introduce additional methods for evaluating and visualizing pipeline outputs.

The `found` library provides multiple entry-points for the HiDDEN pipeline tailored to different use-cases: `found.HiDDEN` for standard use, `found.HiDDENt` for automatic hyperparameter selection, `found.HiDDENg` for automatic splitting by a factor of interest, and `found.HiDDENgt` for combined splitting and hyperparameter selection. All of these high-level entry points accept an `anndata.AnnData` object (which is the de facto standard for -omics work in Python) as input, allowing seamless integration into existing workflows (fig. 1b) [4].

To facilitate evaluation of HiDDEN outputs, `found` also provides a set of utilities used to plot pipeline outputs via the `found.pl` module (fig. 1c). These support evaluating outputs of both `found.{HiDDEN,HiDDENg}` and `found.{HiDDENt,HiDDENgt}`.

Finally, `found` adopts a design based on inversion-of-control software engineering principles, where pipeline steps declare their own data dependencies via argument names rather than relying on fixed inputs. This enables flexible composition of pipeline methods and supports integration of additional data modalities.

Given the widespread use of the R programming language in the -omics analysis space, we also provide `found` as an R package [5]. To ensure consistency between implementations, the R package was designed to utilize the Python library as a back-end, minimizing divergence in functionality and behavior. Mirroring the design of the Python API, `HiDDEN{,g,t,gt}` entry-points are provided as `S7` generics, with methods implemented for both `SingleCellExperiment` and `Seurat` formats (fig. 1c) [6], [7], [8]. Moreover, a `to_step` adapter is provided, allowing for R functions to be used in pipelines where other steps are implemented in Python. This adapter follows the conventions of the Python workflow, as functions are expected to declare their data dependencies via argument names. Given known issues with maintenance difficulty of alternate language implementations (e.g. behavior and/or feature set drift, introduction of bugs during translation, etc.), specific consideration was made to minimize redundancy between the `found` Python and R libraries. Instead of reimplementing Python routines in R, the R `found` package relies maximally on the Python implementation for function, with R code written effectively only for R to- and from-Python format conversions. While this adds a runtime and memory overhead to R `found` use, as time and memory is spent redundantly copying data structures across formats, internal use has shown that within-pipeline computation dominates over these when assessing performance (supplemental fig. 3).

Comprehensive documentation is provided for both Python and R packages (using `sphinx` and `pkgdown` respectively), available publicly at https://goeva-lab.github.io/found/py and https://goeva-lab.github.io/found/R, and is automatically built via GitHub workflows on pushes to the remote main branch [9], [10]. This includes complete public API documentation for each package as well as multiple guides to demonstrate standard API use. For ease of deployment across computational environments, Docker images are built on pushes to main (containing preinstalled Python and R `found` packages), distributed via the GitHub Container Registry at https://ghcr.io/goeva-lab/found. To guarantee validity of Python and R code, automated code quality/correctness checks are conducted on all pushes to the GitHub remote.

### HiDDEN improves sensitivity of downstream analyses

As a representative example, we assessed the utility of HiDDEN outputs on a recently released dataset of IL-15 stimulated PBMCs (with PBS-exposed cells as control) [11]. We evaluated HiDDEN outputs using two approaches to identify differentially expressed genes (DEGs) using negative binomial models [12]. First, to assess whether 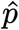 captures biological signal, we conducted differential gene expression analysis using 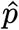 values as model inputs, yielding numerous significantly regulated genes across multiple cell types (fig. 1e). Second, we used 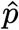 to remove “unaffected” cells from the case group prior to cross-condition DEG analysis of “pseudobulk” sample and cell type aggregates (fig. 1e). This filtering step enabled by our binary refined labels increased the number of DEGs when conducting cell type-specific analysis in this fashion (fig. 1e). These results indicate that HiDDEN-derived per-cell perturbation estimates enhance signal detection in downstream analyses.

### found benchmarking

To evaluate `found` and characterize the behavior of HiDDEN in practice, we performed extensive benchmarking. We systematically assessed the effects of five parameters across ten datasets: preprocessing and embedding method, continuous scoring/regression method, binarization method, *k* selection, and cell identity grouping, where the final parameter determines if HiDDEN is run jointly across all cells or separately for each cell type.

### HiDDEN is sensitive to hyperparameters in a data-dependent manner

Analysis of 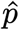 reveals strong sensitivity to the underlying regression method utilized (supplemental fig. 1). We find that logistic regression is well suited for disease continuum analysis, while random forest and support vector machine (SVM) outputs are ill-suited, each for different reasons. Random forest outputs tend to overfit on training data, returning nearly single-mass probability distributions concentrated at 0 for control and 1 for case cells (supplemental fig. 1). This is concordant with prior results analyzing random forest classifiers, which are known to show very high discrimination rates on training data [13]. This makes random forest outputs ill-suited for extracting a disease continuum signal given the task design and hyperparameters. On the other hand, SVM outputs, which are distances from the class-separating-hyperplane rather than probabilities, are generally dense around zero, yielding little to no ability to extract disease continuum signal (supplemental fig. 1). This can be explained by the optimization target of SVMs being solely correct class separation, with no incentive to increase confidence once a satisfying hyperplane is produced. This stands in contrast to logistic regression, where cross-entropy loss penalizes correct but low confidence predictions. This provides an incentive to generate parameters which yield higher 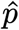 for truly perturbed cells and lower 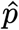 for truly unaffected cells, which in turn increases the ability to distinguish e.g. unaffected case cells as the ones with distinctly lower 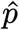.

For other parameters, we observe different levels of sensitivities to individual and interacting parameter changes in a data-dependent manner. Using heuristic-based evaluation, we observed that grouping mode and *k*-selection emerge as major influences, whereas binarization and embedding methods play a subtler role (fig. 2a, supplemental fig. 2). Grouping mode effects appear data-dependent, with splitting by identity leading to generally worse scores on smaller datasets with indications that this trend reverses on larger datasets (fig. 2a). As another example, we observe different optimal *k* values across datasets (fig. 2b), highlighting the need for domain expert evaluation of outputs during HiDDEN application to novel biological questions and corresponding -omics data.

**Figure 2:**
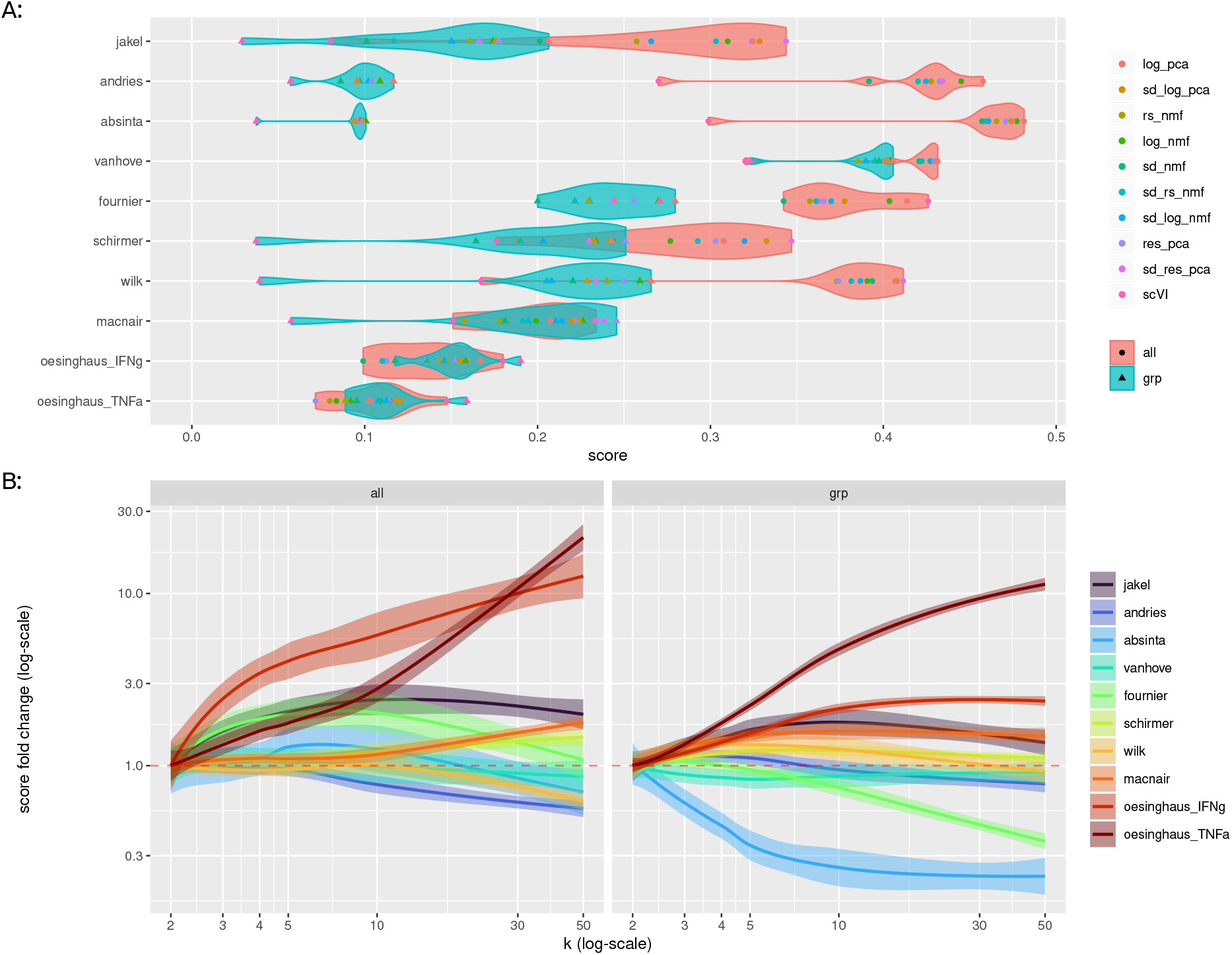
Sensitivity of HiDDEN to modeling choices. A: Null distance scores (x-axis) across embedding methods (point colour), datasets (y-axis), and pipeline grouping configuration (violin fill colour and point shape). Datasets are ordered by size (smallest to largest from top to bottom). Score values are displayed for the corresponding dataset / embedding method / grouping with 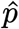 generated via logistic regression, and for the combination of *k* and harmonization application yielding the highest score. B: Inter-group distance scores for 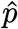 generated via logistic regression from the initial *k* value (*k* = 2) configuration (y-axis, log-scale) across datasets (line/fill colour) and pipeline grouping configuration (left versus right -hand facets) for different *k* values (x-axis, log-scale). Values across binarization methods and embedding methods are grouped for each corresponding configuration. Shaded regions indicate 0.95 confidence intervals around lines which represent smoothed trends (computed via the `loess` function from the R `stats` package, with the x-axis in log-space).

## Conclusions

HiDDEN is a modular framework whose performance depends on modeling choices at each stage of the pipeline. Based on benchmarking across diverse datasets, we provide the following practical recommendations.

For preprocessing and embedding, we recommend shifted-logarithm-transformed and PCA-embedded representations, which are widely used across -omics workflows, exhibit favorable scaling properties (supplemental fig. 3), and show relatively limited sensitivity compared to alternatives. At the same time, `found` supports the use of externally derived embeddings, enabling integration of custom representations, including those designed for cross-sample or cross-species harmonization. For regression, we recommend logistic regression based on the observed limitations of alternative methods. For binarization, given the similar performance of k-means and a two-component Gaussian mixture model (supplemental fig. 2), we recommend k-means due to its on average superior runtime and memory performance profile (supplemental fig. 3).

In terms of pipeline troubleshooting, we recommend prioritizing *k*-selection and grouping, by initially exploring a range of parameter values followed by inspection via scoring metrics and manual assessment via the `found.pl` module.

In summary, we present `found`, a comprehensive implementation of the HiDDEN framework that enables its practical and reproducible use. By exposing HiDDEN as a flexible pipeline with multiple modeling choices and providing diagnostic tools for evaluation, `found` facilitates systematic exploration of modeling choices and parameter settings, thus improving robustness in downstream analyses. In particular, diagnostic scoring and visualization tools enable assessment and validation of inferred perturbation signals.

By combining flexible design with accessible interfaces and comprehensive documentation and usage guides, `found` supports a wide range of users, from domain scientists to method developers. Moreover, it enables efficient exploration of large single-cell - omics datasets across perturbation conditions, facilitating the identification of subtle or heterogeneous perturbation signals that may otherwise remain undetected.

## Methods

### found Python library

The `found` Python library is organized into modules that implement pipeline orchestration, methods, visualization, and hyperparameter tuning:

- `found.adapters` contains code relevant to orchestrating pipeline steps, most notably providing the `found.adapters.Pipeline` class
- `found.find` contains code relevant to the various `HiDDEN` entry-point functions, e.g. `HiDDEN`, `HiDDENg`, `HiDDENt`, and `HiDDENgt`, with data structures from the `anndata` and `pandas` libraries being utilized [4], [14]
- `found.methods` contains code relevant to implementation of various pipeline routines, such as dimensionality reduction, regression, binarization, and output evaluation, relying on the `scikit-learn`, `scipy`, and `numpy` libraries for underlying implementations [15], [16], [17]
- `found.pl` contains code relevant to plotting/evaluating pipeline outputs, relying on the `vega-altair` library for plot creation and display [18], [19]
- `found.seed` contains helper functions used for coordination of a seeding variable, notably used by functions in `found.methods` to allow for reproducible pipeline outputs
- `found.tune` contains code relevant to automatic hyperparameter selection (termed “tuning”), providing out-of-the-box implementations for different selection strategies
- `found.types` contains various type definitions reused across the rest of the package

Overall, the library has direct dependencies on only the following libraries: `anndata`, `scikitlearn`, `scipy`, `numpy`, `pandas`, and `vega-altair`, all of which are already commonly used in the field of high dimensional -omics data analysis.

Building documentation for the library requires extra development dependencies, which are:

- `sphinx`, `sphinx-autobuild`, `sphinx-autodoc-typehints`, `pydata-sphinx-theme`: used for generating HTML documentation pages [10], [20], [21], [22]
- `myst-nb`, `myst-parser`, `jupytext`, `ipykernel`, `ipywidgets`: used for rendering guides, as they are written in the `jupytext` format [23], [24], [25], [26], [27]
- `pyLemur`: used within guides to demonstrate extensibility and integration with other packages [28]

The `uv_build` package is utilized as a build system to generate package builds [29].

### found R library

The `found` R package is composed of only a single R source file (`found.R`), which defines generics `HiDDEN`, `HiDDENg`, `HiDDENt`, and `HiDDENgt`. The package has baseline dependencies on only the following packages: `S7` (used to define generics), `reticulate` (used for R-to-python interoperability), as well as `SeuratObject` and `SingleCellExperiment` [6], [7], [8], [30].

Building the documentation for the package requires extra development dependencies, which are:

- `irlba`, `Seurat`, `SeuratData`/`ifnb.SeuratData` - used in guides to demonstrate functionality [31], [32], [33]
- `pkgdown` - used for generating HTML documentation pages [9]
- `devtools`, `lintr`, `pak` - used for package management and maintenance [34], [35], [36]

### IL-15 differential gene expression analysis

To generate the test dataset, cells from the PBS and IL-15 conditions of the data in [11] were selected. HiDDEN was applied to the resulting data (using PBS-exposed cells as controls) in a grouped fashion (e.g. via the `HiDDENg` entry-point), splitting by the `cell_type` metadata column, with a *k* value of 50 (defaults were used for all other pipeline parameters).

Differential gene expression analysis was then conducted using the `glmGamPoi` R library [12].

For continuous score analysis, a model was fit (using the `glmGamPoi::glm_gp` function) on the full count matrix with 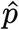 values as the independent variable. The `glmGamPoi::test_de` function was then applied for each cell type separately to generate cell type specific lists of genes significantly regulated along 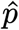.

For discretized label analysis, aggregated matrices were generated from the cell by gene count matrix by pseudobulking across sample, condition, and cell type using the `glmGamPoi::pseudobulk` function. Two such aggregate matrices were generated, one from the full dataset, and another by first removing case cells rendered unaffected by the HiDDEN discretization labels. Resulting matrices were used to fit `glmGamPoi` models, with cell type and condition metadata being provided as independent variables (again using `glmGamPoi::glm_gp`). Cell type specific differentially expressed genes were then extracted between conditions by using the `glmGamPoi::test_de` function as described above.

### Benchmarking

#### Runtime profile extraction

Two metrics were extracted to assess method runtime profiles: peak memory use and runtime. Runtime for each method was extracted using the Python standard library `cProfile` module, by running method-relevant functions using `cprofile.Profile.run_call`, then extracting the `total_tt` field from the derived `pstats.StatsProfile` object [37]. Peak memory use was extracted using the Python `memray` library, by running method-relevant functions under the `memray.Tracker` context, then extracting the `peak_memory` field from the archive-derived `memray.Metadata` object [38].

#### Datasets

GSE and GSM codes below refer to accession numbers from the National Center for Biotechnology Information (NCBI) Gene Expression Omnibus (GEO) archive [39], [40]. Zenodo records refer to accession numbers from the Zenodo archive [41].

Benchmarking was conducted using the following datasets:

- `absinta`: sourced from GSE180759 [42]
- `andries`: sourced from GSM7336928 and GSM7336929 [43]
- `fournier`: sourced from GSM5973615, GSM5973616, GSM5973617, GSM5973620, GSM5973621, and GSM5973622 [44]
- `jakel`: sourced from GSE118257 [45]
- `macnair`: sourced from Zenodo record 8338963 [46]
- `oesinghaus_IFNg`, `oesinghaus_TNFa`: both sourced directly from the data made available at https://parse-wget.s3.us-west-2.amazonaws.com/10m/10M_PBMC_12donor_90cytokines_h5ad.zip [11]
- `schirmer`: sourced directly from the UCSC cell browser website (https://cells.ucsc.edu), at the `/ms/rawMatrix.zip` endpoint [47], [48]
- `vanhove`: sourced directly from the Brain Immune Atlas website (https://www.brainimmuneatlas.org/), at the `/data_files/toDownload/filtered_gene_bc_matrices_mex_irf8_fullAggr.zip` endpoint
- `wilk`: sourced from the CELLxGENE website, as made available at https://datasets.cellxgene.cziscience.com/89c999bd-2ba9-4281-9d22-4261347c5c78.h5ad [49], [50]

The following table outlines the datasets in question

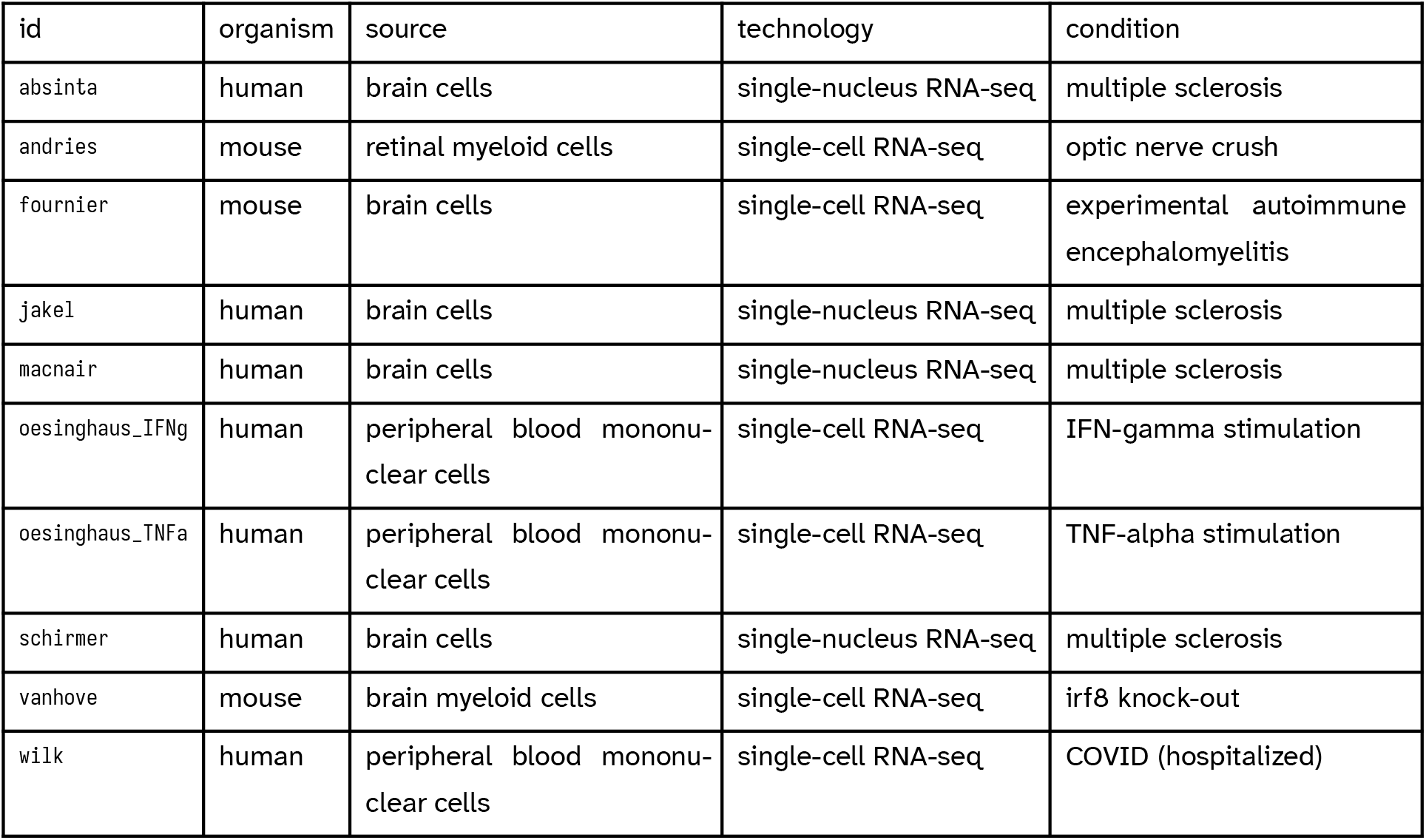

Further information regarding filtering steps and extraction of cell identity, batch, and condition per-observation metadata can be found within the definition of the `DATA_CONF` object defined within the benchmarking script made available with this publication. This script utilized the `cappa` library for argument parsing [51].

#### Preprocessing methods

##### Shifted-logarithm -based transform

A shifted logarithm based transform was implemented as described in [2]. Mathematically, this transform can be defined as follows:

- *X*_*ij*_ refers to the value of the *i*-th cell and the *j*-th gene in the (*i, j*) position in the count matrix
- *θ* refers to the overdispersion factor, here set to 0.05
- log refers to the natural logarithm function
- *S*_*i*_ refers to the sum of all counts for the *i*-th cell
- *S*_*h*_ refers to the geometric mean of *S*_*i*_ for all cells

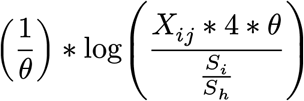

##### Analytic Pearson residual -based transform

An analytic Pearson residual-based transform was implemented as defined in equations (3) and (4) of [52].

#### Embedding methods

##### Matrix factorization

PCA and NMF-based embedding was implemented using the functionality provided by the `sklearn.decomposition.PCA` and `sklearn.decomposition.NMF` classes from the `scikit-learn` package [15]. For PCA, the ARPACK solver was utilized, whereas the coordinate descent method was utilized for NMF [53], [54].

##### Harmony adjustment

To control for the presence of possible biologically-irrelevant batch effects within datasets, for each matrix factorization based embedding method, an additional variant was generated were resulting outputs were adjusted using per-dataset batch factors. To run the Harmony adjustment, the R `harmony` package was used, called from Python via the `rpy2` bridging library [3], [55].

##### scVI embedding

scVI based embedding was conducted on each dataset using the `scvi-tools` python library, by constructing an `scvi.model.SCVI` object, calling the `train` method, and then using the outputs of the `get_latent_representation` method [56].

##### Complete method list

The following table outlines the total number of embedding methods benchmarked, each of which was run for *k* values *k* = 2, *k* = 3, *k* = 4, *k* = 5, *k* = 10, *k* = 30, and *k* = 50:

| id | preprocessing | feature-wise variance scaling | embedding method |
| --- | --- | --- | --- |
| log_pca | shifted-logarithm-based transform | no | PCA |
| res_pca | analytic Pearson residual transform | no | PCA |
| sd_log_pca | shifted-logarithm-based transform | yes | PCA |
| sd_res_pca | analytic Pearson residual transform | yes | PCA |
| sd_nmf | none | yes | NMF |
| rs_nmf | scaling by observation-wise sum | no | NMF |
| log_nmf | shifted-logarithm-based transform | no | NMF |
| sd_rs_nmf | scaling by observation-wise sum | yes | NMF |
| sd_log_nmf | shifted-logarithm-based transform | yes | NMF |
| log_pca_harmony | shifted-logarithm-based transform | no | PCA + harmony |
| res_pca_harmony | analytic Pearson residual transform | no | PCA + harmony |
| sd_log_pca_harmony | shifted-logarithm-based transform | yes | PCA + harmony |
| sd_res_pca_harmony | analytic Pearson residual transform | yes | PCA + harmony |
| sd_nmf_harmony | none | yes | NMF + harmony |
| rs_nmf_harmony | scaling by observation-wise sum | no | NMF + harmony |
| log_nmf_harmony | shifted-logarithm-based transform | no | NMF + harmony |
| sd_rs_nmf_harmony | scaling by observation-wise sum | yes | NMF + harmony |
| sd_log_nmf_harmony | shifted-logarithm-based transform | yes | NMF + harmony |
| scVI | none | no | scVI |

##### Scoring/regression methods

Logistic regression, random forest classifiers, and support vector machines were implemented using functionality provided by the `sklearn.linear_model.LogisticRegression`, `sklearn.ensemble.RandomForestClassifier`, and `sklearn.svm.LinearSVC` classes respectively from the `scikit-learn` package. No l1 or l2 regularization was applied for logistic model fitting, and the L-BFGS solver was utilized [15]. For support vector machines, an l1 penalty was specified and the regularization parameter `C` was set to 1. For all other parameters, default `scikit-learn` values were utilized.

##### Binarization/clustering methods

K-means and Gaussian mixture modeling-based clustering methods were implemented using functionality provided by the `sklearn.mixture.GaussianMixture` and `sklearn.cluster.KMeans` classes provided by the `scikit-learn` package [15]. In both cases, the total number of desired clusters was set to 2. For all other parameters, default `scikitlearn` values were utilized.

##### Heuristic-based evaluation of HiDDEN outputs

Evaluation of HiDDEN outputs was done according to two different heuristics. For all below steps, Earth Mover’s distances (also known as Wasserstein distances) were computed using the `wasserstein` function provided by the R `approxOT` package, with the `p` argument set to 1 (representing the power of the distance), and the `method` argument set to `univariate` [57].

The metric referred to as inter-group distance represents the difference between the Earth Mover’s distances for two different terms. The left-hand side is the distance between the distributions of 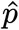 for case cells labeled as affected and case cells labeled as unaffected. The right-hand side is the distance between the distributions of 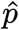 for control cells and case cells labeled as unaffected.

The metric referred to as null distance represents the Earth Mover’s distance between the distribution of 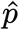 for all analyzed cells and the distribution of 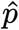 for a simulated null distribution of 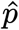. A null distribution is generated by running the regression step with unchanged source data embeddings but with condition labels randomly permuted.

## Availability of data and materials

Source code for the `found` library is available at https://github.com/goeva-lab/found. Source code for the benchmarking script and associated analyses is available at https://github.com/goeva-lab/afanasiev-2026-found_lib.

## Ethics declarations

No human subjects used.

## Large language model (LLM) use

EA did not use LLM-based technologies for any work in or associated with this manuscript, be it in code or manuscript components.

## Authors’ contribution

AG conceived the study, supervised the project, and secured funding. EA designed and implemented the Python and R `found` packages, along with associated workflows, documentation, and infrastructure. AG and EA designed the benchmarking framework. EA implemented and executed the benchmarking framework and analyzed the results. EA drafted the manuscript. All authors reviewed and approved the manuscript and agree to be accountable for their contributions.

## Acknowledgments

The authors would like to thank Dr. Delaram Pouyabahar, Arhan Rupani, Yuxi Zhu, Marcia Jude and Sam Salitra for their help with testing the `found` library. We would also like to thank the Digital Research Alliance of Canada (https://alliancecan.ca), Compute Ontario (https://www.computeontario.ca), and the SHARCNET consortium (https://www.sharcnet.ca) for enabling the work presented here via our use of the Nibi cluster.

## Supplemental Figure 1

**Supplemental Figure 1:**
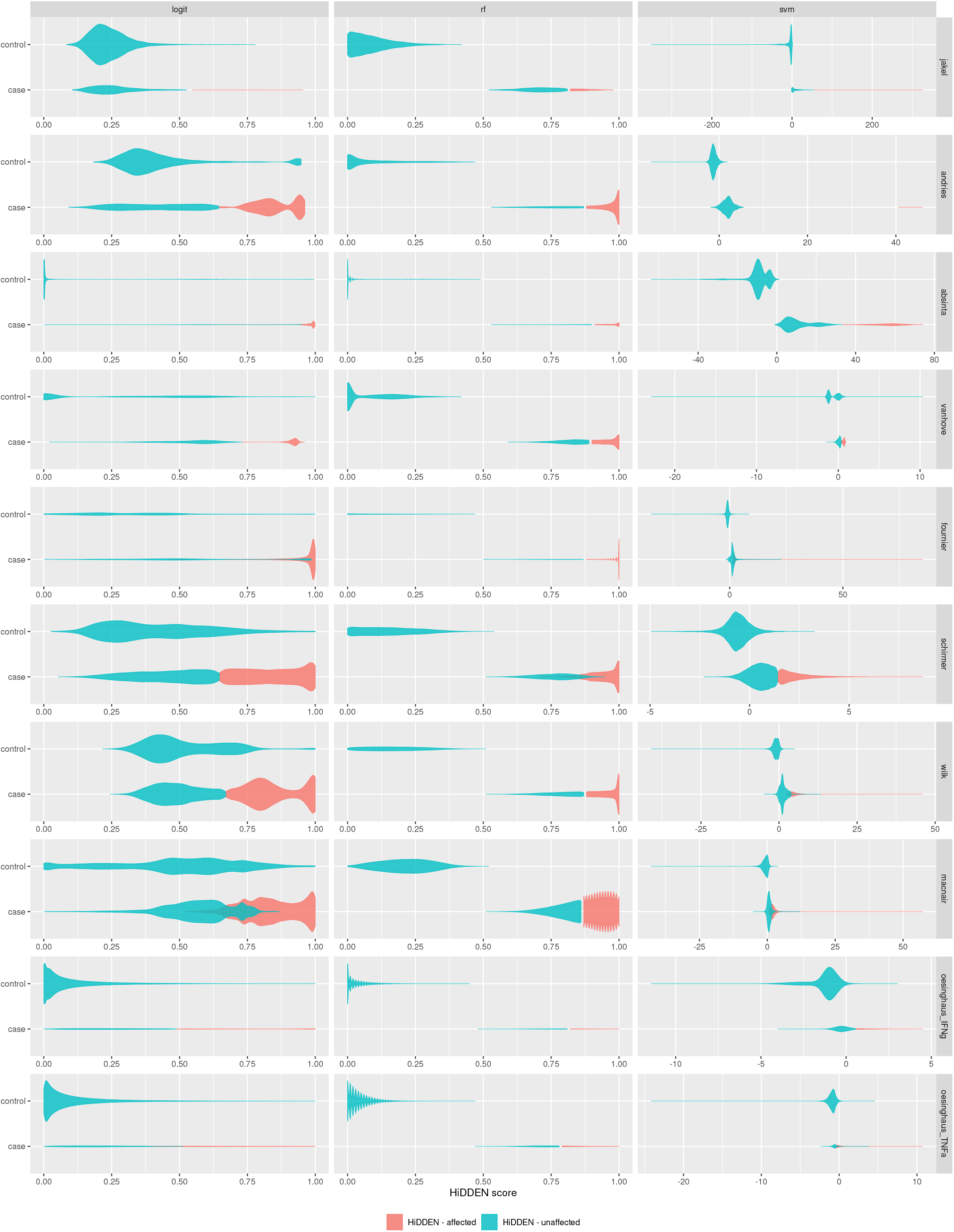
Effect of regression method on inferred perturbation scores. Violin plots of 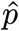 (x-axis) in control and case cells (y-axis), stratified by HiDDEN-refined labels (violin fill colour), datasets (facet rows, ordered by dataset size, smallest to largest from top to bottom) and regression method (facet columns, in left-to-right order: logistic regression, random forest, and support vector machines). Values are shown for 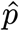 based off of embeddings generated using the `sd_log_pca method`, with *k*, post-hoc harmonization, and pipeline grouping configuration selection based on best inter-group distance scores.

**Supplemental Figure 2:**
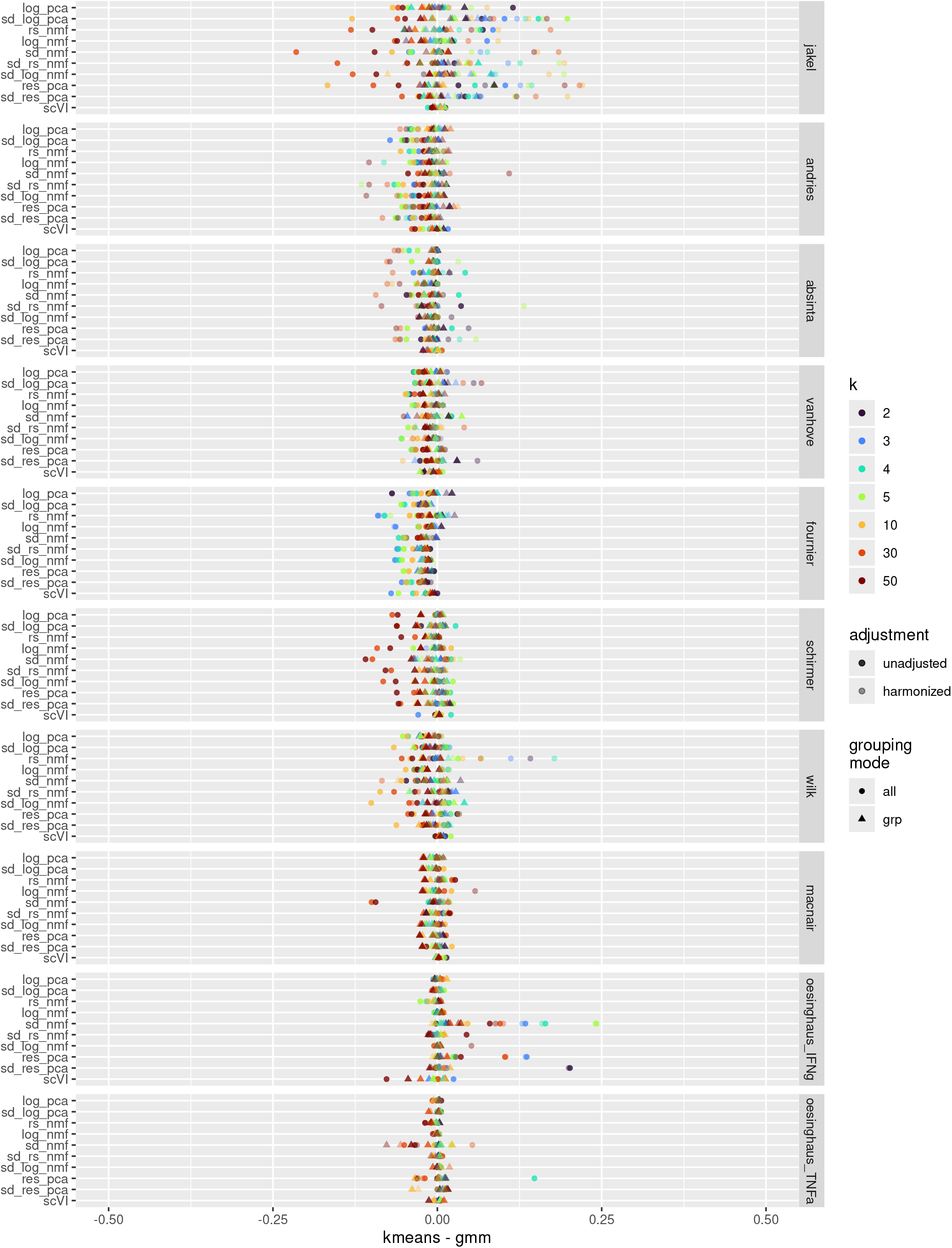
Comparison of binarization methods. Dot plots showing difference in inter-group distance scores between k-means and Gaussian mixture model-based binarization across datasets (facet rows, ordered by dataset size, smallest highest, largest lowest), embedding methods (y-axis), harmonization (point opacity), *k* value (point colour), and pipeline grouping configuration (point shape). Movement to the left on the x-axis represents a larger score using Gaussian mixture modeling, whereas movement to the right represents a larger score using k-means clustering.

**Supplemental Figure 3:**
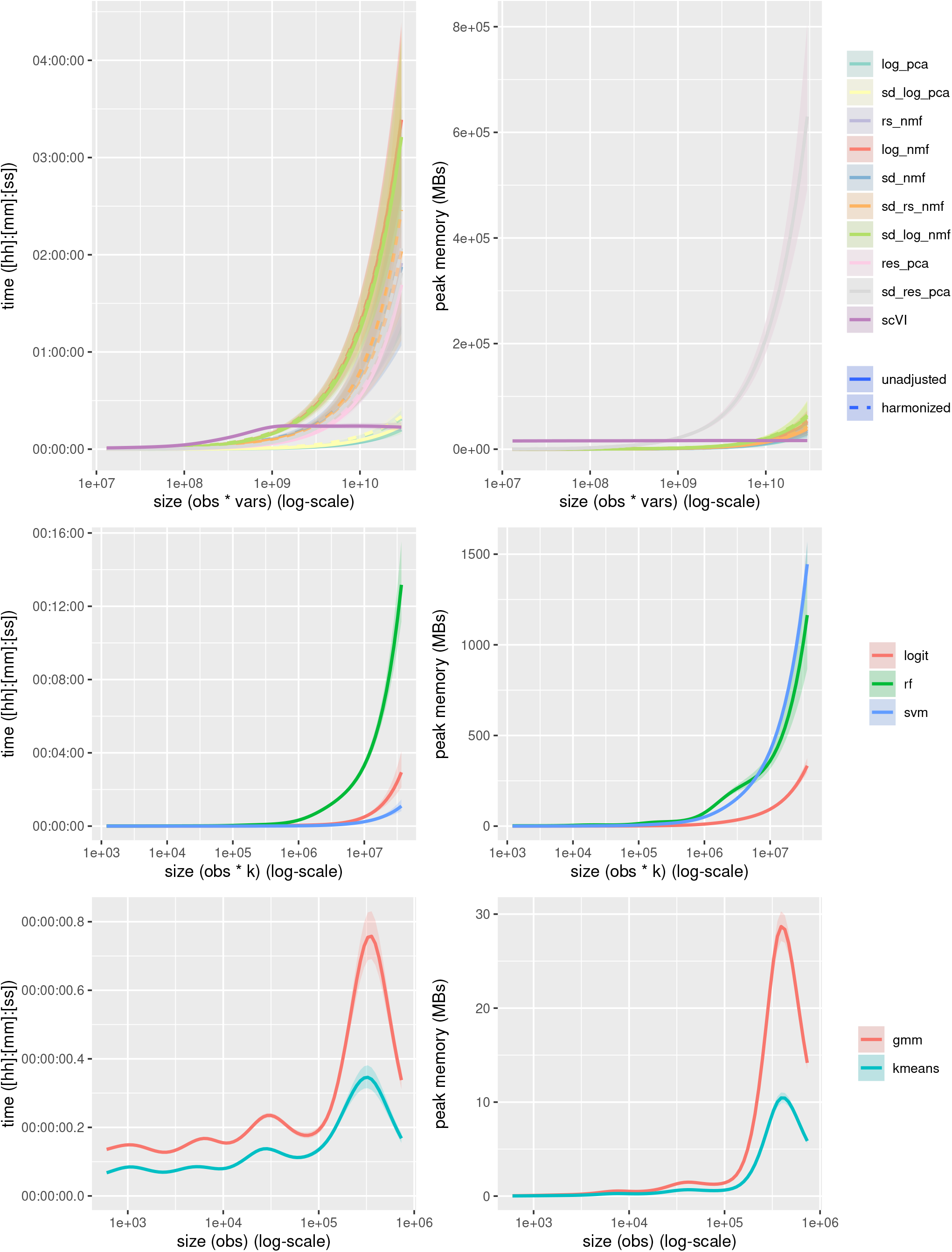
Runtime and memory usage across pipeline components. Runtime and peak memory usage (y-axis, runtime on the left and peak memory use on the right) for embedding, regression, and binarization methods. Lines represent smoothed trends; shaded regions indicate 0.95 confidence intervals computed via the `gam` function from the R `mgcv` package, with y-axis data projected to log-space to enforce the non-negativity constraint of the underlying data [58].

